# Macro-scale models for fluid flow in tumour tissues: impact of microstructure properties

**DOI:** 10.1101/2020.07.02.180026

**Authors:** Cristina Vaghi, Raphaelle Fanciullino, Sebastien Benzekry, Clair Poignard

**Affiliations:** Team MONC, Inria, Institut de Mathématiques de Bordeaux, CNRS, Bordeaux INP, Univ. Bordeaux, France; SMARTc, CRCM Inserm UMR1068, CNRS UMR7258, Aix Marseille University, France

**Keywords:** two-scale homogenisation, fluid flow in tumours, interstitial fluid pressure, tumour microscopic structure

## Abstract

Understanding the dynamics underlying fluid transport in tumour tissues is of fundamental importance to assess processes of drug delivery. Here, we analyse the impact of the tumour microscopic properties on the macroscopic dynamics of vascular and interstitial fluid flow by using formal asymptotic techniques.

Here, we obtained different macroscopic continuum models that couple vascular and interstitial flows. The homogenization technique allows us to derive two macroscale tissue models of fluid flow that take into account the microscopic structure of the vessels and the interstitial tissue. Different regimes were derived according to the magnitude of the vessel wall permeability and the interstitial hydraulic conductivity. Importantly, we provide an analysis of the properties of the models and show the link between them. Numerical simulations were eventually performed to test the models and to investigate the impact of the microstructure on the fluid transport.

Future applications of our models include their calibration with real imaging data to investigate the impact of the tumour microenvironment on drug delivery.

## 1 Introduction

Interstitial and capillary fluids are strongly connected in malignant tissues and are mainly involved in the transport of molecules in tumours. When drugs are intravenously injected, they have to overcome several barriers, including vascular transport, transvascular transfer, interstitial transport and finally cellular uptake [1]. The biological and physicochemical properties of the tumour microenvironment play a significant role in the drug delivery process [2]. The geometrical microstructure of the tumour also has an important impact on the fluid flow [3].

Neoplastic tissues are highly heterogeneous. They are generally characterized by [4] accumulated solid stress [5], abnormal blood vessels network [6], elevated interstitial fluid pressure (IFP) [7], that almost equals the microvessel pressure (MVP) and dense interstitial structure [8]. These traits, that distinguish tumour tissues from normal ones, cause barriers to drug delivery [1]. The heterogeneous spatial distribution of tumour vessels and poor lymphatic drainage impair a uniform delivery of therapeutic agents in tumours. Blood vessels are unevenly distributed, leaving avascular spaces. Moreover, their walls are leaky and hyperpermeable in some places while not in other [9]. Blood flow velocity is also compromised by the elevated viscous and geometrical resistance offered by the tumour vasculature [3]. Finally, the lack of an efficient lymphatic network inside the tumour coupled with leaky tumour vessels leads to a high IFP [10] almost equal to the microvascular pressure [7]. Due to elevated IFP, the tumour interstitium is characterized by no pressure gradient [11, 12].

Several mathematical models have been developed during the last decades to investigate the features of fluid transport in the tumour microenvironment. The porous medium theory has been employed to model interstitial fluid flow (IFF) relying on Darcy’s law and using average field variables defined over the whole tissue [13, 14]. Fluid transport through the blood vessels has been exploited in both discrete and continuous manners, including spatial and temporal variations. In either discrete and continuous models, the IFF and microvascular fluid (MVF) are usually coupled by Starling’s law [15], that describes the fluid filtration through the highly permeable vessels walls. Microscopic models of the flow patterns around an individual capillary and a network of blood vessels have been introduced relying on the Krogh cylinder model [16, 17, 18]. Poiseuille’s law can be considered to describe the blood flow in a cylindrical domain [19, 20, 21]. Furthermore, Navier-Stokes equations have been adopted to model the spatio-temporal variations in blood flow [18, 14]. More detailed biophysical models have been developed to take into account the more realistic heterogeneity of the tumour vasculature [22]. Welter et al [23] introduced an exhaustive biophysical model the incorporates tumor growth, vascular network (including arteries and veins), angiogenesis, vascular remodeling, porous medium description for the extracellular matrix (ECM) and interstitial fluid, interstitial fluid pressure and velocity and chemical entities (such as oxygen, nutrients, drugs). On the other hand, continuous models based on mixture theory have been exploited to describe interstitial and vascular fluid flow, assuming that the two phases are present at each point of the tumour [24]. Multiscale models have further been employed to investigate the coupling between tumor growth, angiogenesis, vascular remodelling and fluid transport [25] and the impact of collagen microstructure on interstitial fluid flow [26]. Imaging data have been integrated to both continuum and discrete models to quantify the effect of the heterogeneity on the fluid transport [27, 28].

The increasing amount of imaging data makes it possible to recover vascular networks in details. However, solving discrete models on the entire vessel tree might be computationally expensive. The formal two-scale homogenization technique allows to take into account microscopic features on the macroscopic dynamic of fluid flow. Two-scale asymptotic expansion has been previously applied to fluid and drug transport in tumours. A system of Darcy’s equations has been derived in [29] to couple interstitial and vascular fluid flows in malignant tissues assuming a periodic medium. A higher complexity has been taken into account in [30], with the introduction of rheological effects in the blood flow and of local heterogeneity. A generalization of homogenized modelling for vascularized poroelastic materials has also been presented [31, 32]. More recently, higher complexity has been added to the homogenized models [33] considering three length scales for the vessel network (i.e., arteriole, venule and capillary scales).

The main novelty of this work is the study of the impact of the tumour microscopic properties on the global fluid dynamic. First, we describe a system of partial differential equations coupling interstitial, transvascular and capillary flows at the microscopic scale. While the interstitium and the capillary walls are assumed to be porous media where the fluid is governed by Darcy’s law, the blood in the capillaries is considered a Newtonian fluid described by the Stokes equation. As the thickness of the capillary walls tends to zero, an asymptotic analysis similar to [34, 35] enables us to derive a Starling’s law-type equation across the vessel wall for the transvascular transport of fluid flow. Then, we perform a two-scale analysis under periodic assumption [36] to derive formally effective macroscale tissue models of fluid flow for 3 asymptotic regimes depending on the magnitude of the permeability of the vessel wall and of the interstitial hydraulic conductivity. These models combine the effects of the interstitial compartment and the capillaries at the microscale, providing thus a link between the microstructure and the macroscopic fluid flow. Moreover, we compare the different asymptotic regimes with some models given in literature (namely, [2], [29], [30]) and show the links between the different models. In particular, we show that for the model initially derived by Shipley and Chapman [29] the difference between the capillary and the interstitium pressures decays exponentially fast from the boundary, making thus a link with the Baxter and Jain model [2]. Furthermore, we present the mathematical and numerical analysis on the homogenised tensors in order to assess their properties according to the geometric microstructure. Eventually, numerical simulations on the macroscopic models are performed and the results are compared to the literature.

This approach can be applied to study the impact of the tumour microscopic characteristics on drug delivery. Imaging data can provide the tissue microstructure that can be integrated in the homogenised model. This modelling technique prevents the resolution of the original micro-scale model that might be unfeasible as it requires the discretisation of the entire vessel network and porous medium. Moreover, the heterogeneities of malignant tissues can be taken into account by considering the spatial variability of the micro-vessel features at the macroscopic scale.

## Main results

First, we motivated the interface conditions between the interstitial compartment and the capillaries of the micro-scale model using an asymptotic expansion technique. We obtained a Starling’s law type equation to describe the flux through the vessel walls. Moreover, we considered the Joseph-Beavers-Saffman slip condition at the boundary between the capillary lumen and the vessel wall. This condition states that the slip velocity along the vessel wall is proportional to the shear stress [37].

Then, using the two-scale asymptotic homogenisation, we assumed that a generic variable *v*^*ε*^(**x**), as function of the macroscopic spatial variable **x** and of the microscopic parameter *ε*, could be approximated as

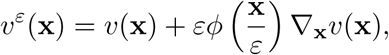

where *v*(**x**) is the macroscopic variable and 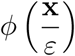 is the corrector that takes into account oscillations at the microscopic scale. Using this approach, we derived three different macroscale models to describe the fluid transport in tumour tissues according to the magnitude of the permeability of the vessel walls and of the interstitial hydraulic conductivity. In particular, the following regimens were derived for the interstitial fluid pressure *p*_*t*_ and the capillary pressure *p*_*c*_:

- **Model 1**: highly permeable walls and large interstitial hydraulic conductivity

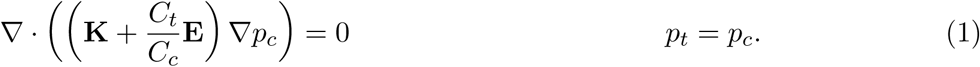
- **Model 2**: weakly permeable walls and large interstitial hydraulic conductivity

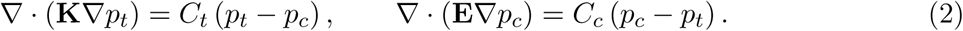
- **Model 3**: weakly permeable walls and small interstitial hydraulic conductivity

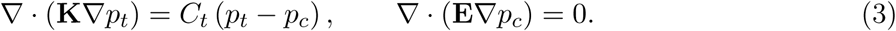

In the three models, **K** and **E** are the permeability tensors of the interstitium and the capillaries, respectively, and *C*_*t*_, *C*_*c*_ are constant parameters that will be defined later on. The definition of these tensors involves the so-called correctors, as usual in homogenisation. It characterises the impact of the microstructure on the effective properties of the tissue at the macroscale. Model (2) has been derived assuming a small magnitude of the capillary permeability and confirms previous results [29, 30].

The interstitial fluid flow **u**_*t*_ and the blood velocity **u**_*c*_ are given in the first two cases by the average quantities

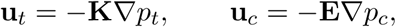

while for the third model they are given by

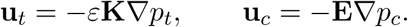

Boundary conditions must be added in order to guarantee well-posedness. Dirichlet-Dirichlet boundary conditions can be imposed to *p*_*t*_ and *p*_*c*_ if the pressures in the sourranding tissue are known. Mixed Dirichlet and Neumann boundary conditions can be chosen to ensure the continuity of **u**_*t*_ or **u**_*c*_.

## 2 Microscale model of fluid transport i n tumours

At the microscale, the domain Ω ∈ ℝ^*N*^ (with *N* = 2, 3) is the medium that consists of the interstitium Ω_*t*_ and the capillary region Ω_*c*_ (Fig. 1). The interface between the capillary and the vessel wall and the one between the interstitium and the vessel wall are denoted respectively by Γ = ∂Ω_*c*_ ∩ ∂Ω_*m*_. In the two regions, the fluid flow is assumed to be incompressible.

**Figure 1.**
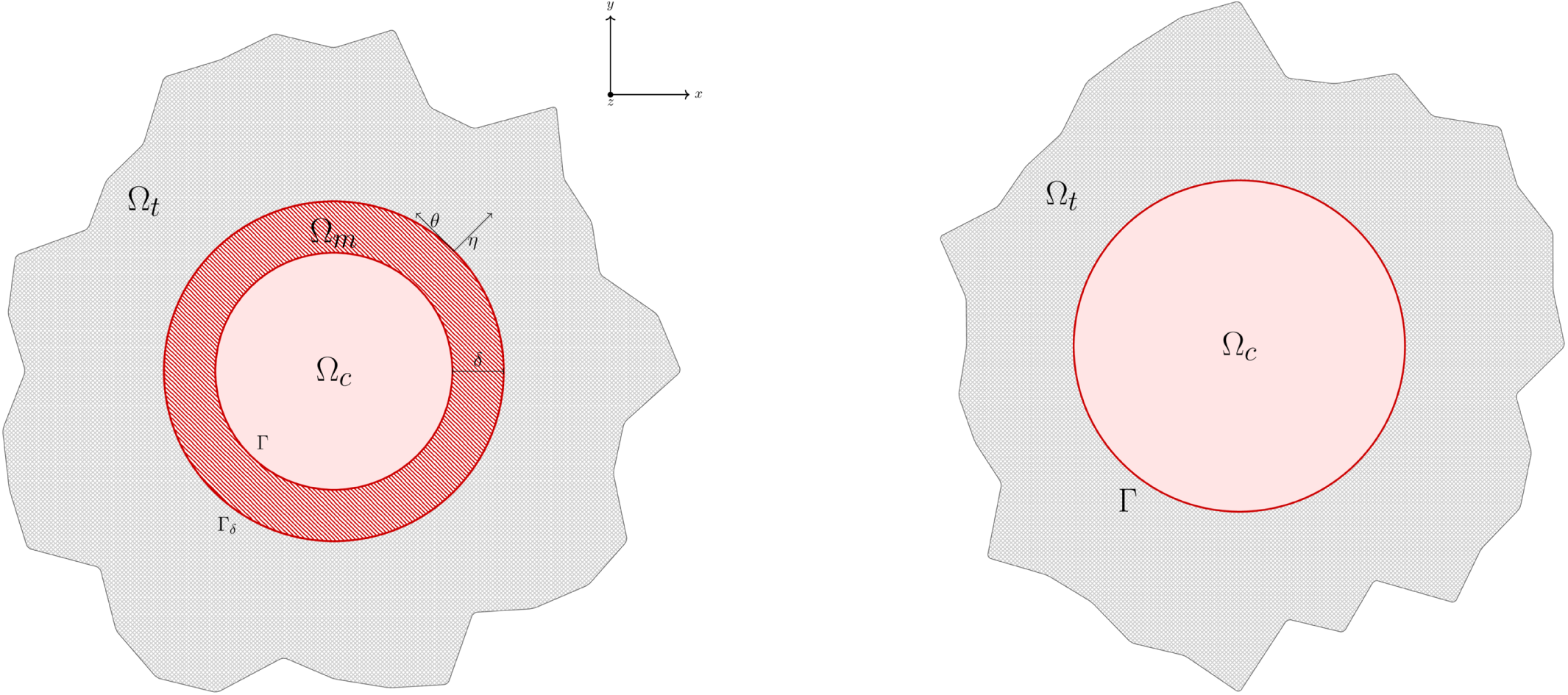
*Left:* schematic of the domain considered to compute the interface conditions between the capillaries and the interstitium. Ω_*m*_ denotes the vessel wall region, Γ_*δ*_ is the interface between the vessel wall and the interstitium. The transmission conditions are derived as *δ*→ 0 using asymptotic expansion. *Right:* schematic representation of the domain of the microscopic model: section of the capillary in the sourranding tissue.

The interstitium - composed by the cells and the extracellular matrix and collagen - is modeled as an isotropic porous medium, where the velocity **u**_*t*_ and pressure *p*_*t*_ follow the Darcy’s law:

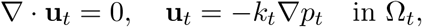

where *k*_*t*_ is the hydraulic conductivity in the interstitium.

In the capillaries, we assume that the fluid is Newtonian with a constant viscosity *µ*. Neglecting the inertial effects and under the assumption of a laminar flow, we end up with the Stokes equation for the description of the vessel velocity **u**_*c*_ and pressure *p*_*c*_

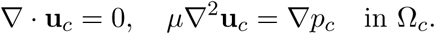

### Interface conditions

Along the vessel wall Γ, interface conditions couple the different equations in the two domains. We performed an asymptotic expansion in order to derive suitable interface conditions between the capillaries and the interstitial compartment as shown in Supplementary Information S1. We considered the vessel membrane as a thin and weakly porous medium and obtained the following interface conditions from the thin layer model, getting similar results as [29, 30]:

1. Continuity of the normal velocity:

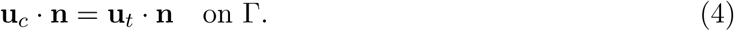 This condition guarantees the continuity of mass through the interface and it is a natural choice since the fluid is assumed to be incompressible in the two regions.
2. Starling ‘s law-type equation, that is the most widely used equation in literature to model flux transport across the vessel wall [12, 19]:

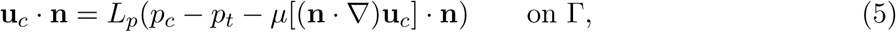

where *L*_*p*_ is the vascular permeability. The difference between the osmotic pressures was considered negligible compared to the interstitial fluid pressure difference in tumours [13, 38]. Moreover, the viscous term in equation (5) is usually neglected but it is necessary to guarantee the well-posedness of the problem and does not change the physical meaning since it is based on the balance of the normal forces [30].
3. Beavers-Joseph-Saffmann condition on the tangential component of the capillary velocity at the boundary with a porous medium:

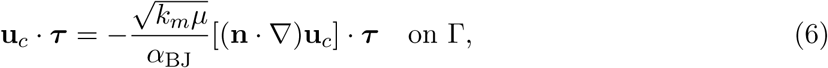

where *α*_BJ_ is a constant depending on the properties of the interface and *k*_*m*_ is the hydraulic conductivity of the vessel walls. This condition comes from the experimental evidence shown by Beavers and Joseph [37] who observed that the slip velocity along Γ was proportional to the shear stress along Γ. Equation of the form (6) was derived by Saffmann using a statistical approach and the Brinkman approximation for non-homogeneous porous medium [39].

### Non-dimensionalization

We perform a dimensional analysis in order to understand the relative amplitude of the different parameters involved. We rescale our fields as follows:

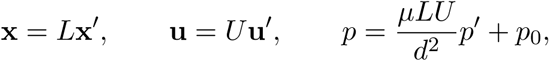

where *L* is the characteristic domain length, *d* is the mean intercapillary distance and *U* is a characteristic velocity. The non-dimensional fluid transport problem reads (neglecting the primes for the sake of simplicity)

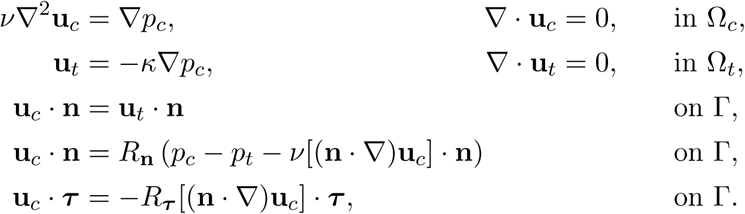

where

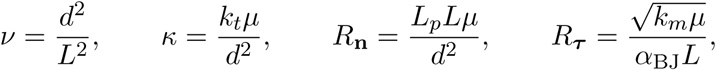

are dimensionless quantities.

## 3 Continuum macroscale models using two scale asymptotic analysis

This section is devoted to the derivation of a continuum macro-scale model using the two scale asymptotic expansion method [40, 41, 36]. We assume that *d* is the mean inter-capillary distance and *L* is the tissue characteristic length such that *ε* = *d/L* ≪ 1. We denote by *Y* the reference periodic cell that is contained in [0, 1]^*N*^ and is composed by the interstitium *Y*_*t*_ and the capillaries *Y*_*c*_, i.e. *Y* = *Y*_*c*_ ∪ *Y*_*t*_ and the interface Γ_*Y*_ = ∂*Y*_*c*_ ∩ 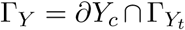. We assume that the interface is entirely contained in *Y*, i.e. Γ_*Y*_ ∩ ∂*Y* = ∅. The normal vector **n** to the interface Γ_*Y*_ is directed outward the vascular domain *Y*_*c*_. The total domain Ω is divided periodically in each direction in identical squares 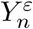 such that

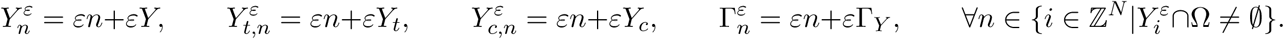

Therefore, the domain Ω is composed of two subdomains 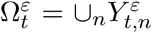 and 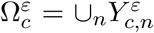 that depend on *ε* and are connected when *N* = 3. The interface between the two subdomains is 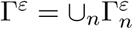. Figure 2 shows a schematic illustration of the periodic domain and of the unitary cell *Y*. According to the multiple scales theory, we introduce a spatial variation decoupling

**Figure 2.**
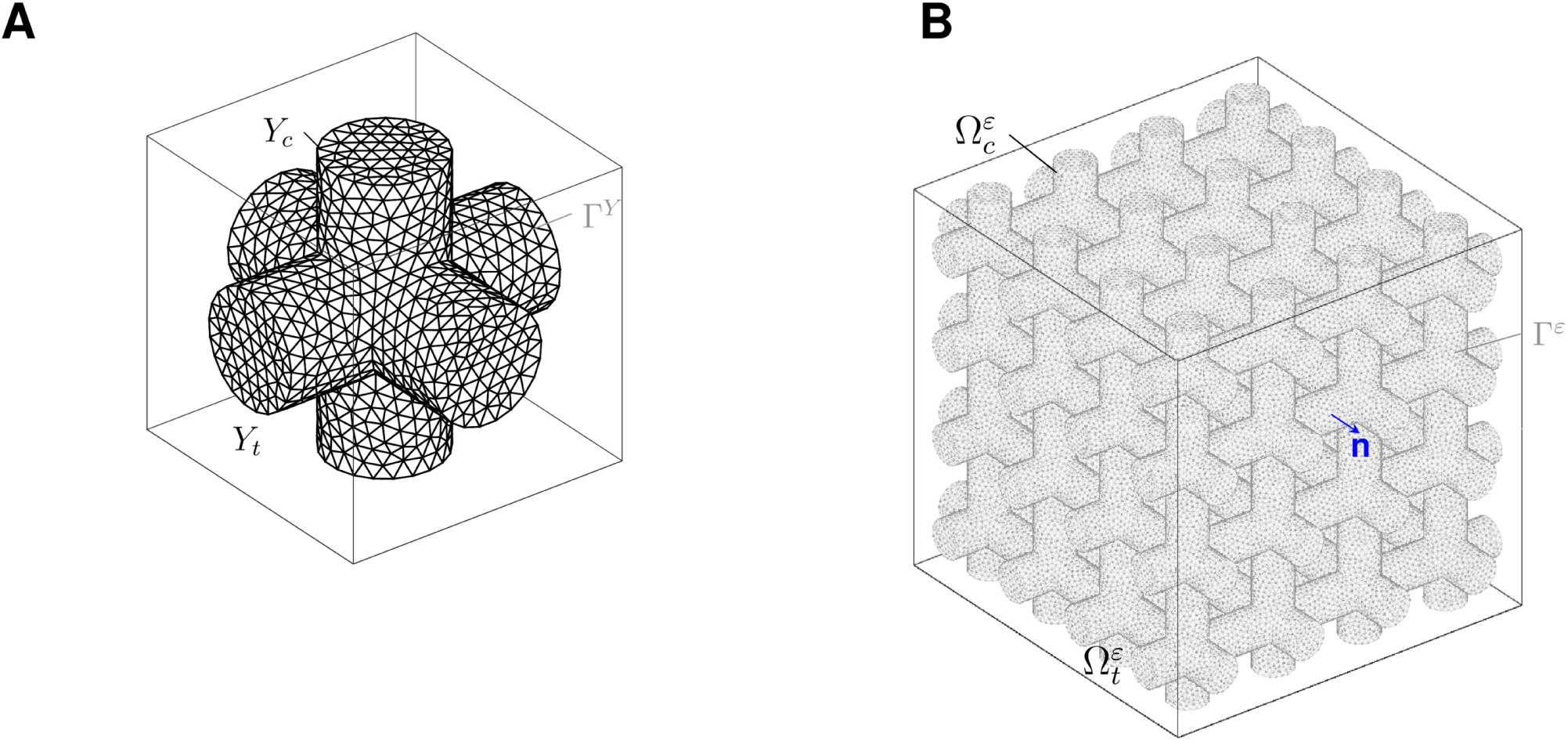
Unitary cell *Y* = [0, 1]^3^ (left) composed by the capillary region *Y*_*c*_ and the interstitial compartment *Y*_*t*_; the interface between the two regions is denoted by Γ_*Y*_. Periodic domain Ω (right): the tumour capillaries 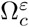 are assumed to be in the tubes, while the outer region corresponds to the interstitial compartment 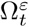; the interface between the two regions is denoted by Γ^*ε*^ and the normal **n** is directed outward the vascular domain.

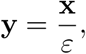

and assume that the macro and the micro spatial variables (**x** and **y**, respectively) are independent. Therefore, any field *g* that we have introduced before (such as **u**_*t*_, **u**_*c*_, *p*_*t*_, *p*_*c*_) depends on *ε* and is assumed to be written following an asymptotic expansion as:

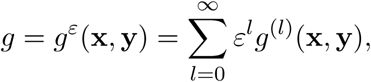

where *g*^(*l*)^(**x, y**) are *Y* -periodic functions.

Assuming that *ν* is of the same scale of *ε*^2^, that *R*_**n**_ is of the order of *ε*^*γ*^ (with *γ* = 0, 1, 2) and that *κ* is of the order of *ε*^*η*^ (with *η* = 0, 1), we rescale the three parameters as 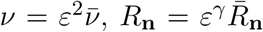 and 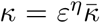 (so that 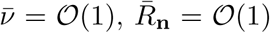 and 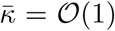. The choice of the rescaling of the parameter *ν* is made in order to avoid trivial solution when *ε* → 0. We analyse the different regimens on varying the exponent *γ*.

We summarize here the main results relative to the leading order and derive formally the homogenized models in the Supporting Information S2. Let us denote by 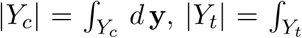dy and 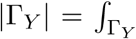 *d s* the measures of the subdomains *Y*_*c*_ and *Y*_*t*_ and of the interface Γ_*Y*_, respectively. We introduce the following two cell problems on the *Y* -periodic cell variables G^*j*^, **W**_*j*_ and P_*j*_ (*j* = 1, …, *N*):

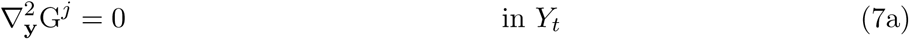

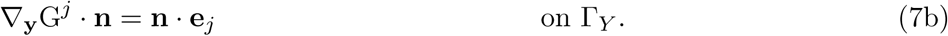

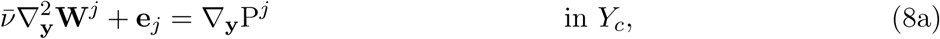

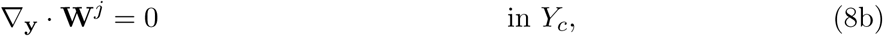

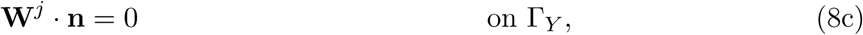

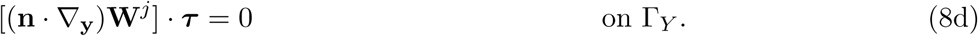

Let us define the following tensors:

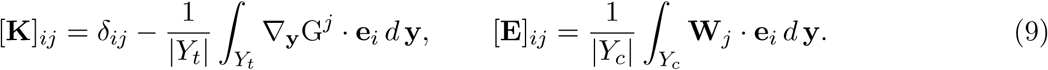

The equations for the leading orders are summarized in Table 1. Boundary conditions need to be added, such as Dirichlet or Neumann conditions.

**Table 1.** Main results of the two-scale asymptotic analysis on varying the order of the parameters *R*_**n**_ and *κ*.

| Case | Convergence | Equation for the leading order in $\Omega$ |
| --- | --- | --- |
| (1) $\gamma = 0, \eta = 0$ | $p_t^\varepsilon(\mathbf{x}, \mathbf{y}) \rightharpoonup p_t^{(0)}(\mathbf{x})$ | $\nabla \cdot \left( \left( \bar{\kappa} \mathbf{K} + \frac{\vert Y_c\vert }{\vert Y_t\vert } \mathbf{E} \right) \nabla p_t^{(0)} \right) = 0$ |
| | $\mathbf{u}_t^\varepsilon(\mathbf{x}, \mathbf{y}) \rightharpoonup \mathbf{u}_t^{(0)}(\mathbf{x}, \mathbf{y})$ | $\langle \mathbf{u}_t^{(0)} \rangle_{Y_t} = -\bar{\kappa} \mathbf{K} \nabla p_t^{(0)}$ |
| | $p_c^\varepsilon(\mathbf{x}, \mathbf{y}) \rightharpoonup p_c^{(0)}(\mathbf{x})$ | $p_c^{(0)} = p_t^{(0)}$ |
| | $\mathbf{u}_c^\varepsilon(\mathbf{x}, \mathbf{y}) \rightharpoonup \mathbf{u}_c^{(0)}(\mathbf{x}, \mathbf{y})$ | $\langle \mathbf{u}_c^{(0)} \rangle_{Y_c} = -\mathbf{E} \nabla p_c^{(0)}$ |
| (2) $\gamma = 1, \eta = 0$ | $p_t^\varepsilon(\mathbf{x}, \mathbf{y}) \rightharpoonup p_t^{(0)}(\mathbf{x})$ | $\nabla \cdot \left( \bar{\kappa} \mathbf{K} \nabla p_t^{(0)} \right) = \frac{\bar{R}_{\mathbf{n}} \vert \Gamma_Y\vert }{\vert Y_t\vert } (p_t^{(0)} - p_c^{(0)})$ |
| | $\mathbf{u}_t^\varepsilon(\mathbf{x}, \mathbf{y}) \rightharpoonup \mathbf{u}_t^{(0)}(\mathbf{x}, \mathbf{y})$ | $\langle \mathbf{u}_t^{(0)} \rangle_{Y_t} = -\bar{\kappa} \mathbf{K} \nabla p_t^{(0)}$ |
| | $p_c^\varepsilon(\mathbf{x}, \mathbf{y}) \rightharpoonup p_c^{(0)}(\mathbf{x})$ | $\nabla \cdot \left( \mathbf{E} \nabla p_c^{(0)} \right) = \frac{\bar{R}_{\mathbf{n}} \vert \Gamma_Y\vert }{\vert Y_c\vert } (p_c^{(0)} - p_t^{(0)})$ |
| | $\mathbf{u}_c^\varepsilon(\mathbf{x}, \mathbf{y}) \rightharpoonup \mathbf{u}_c^{(0)}(\mathbf{x}, \mathbf{y})$ | $\langle \mathbf{u}_c^{(0)} \rangle_{Y_c} = -\mathbf{E} \nabla p_c^{(0)}$ |
| (3) $\gamma = 2, \eta = 1$ | $p_t^\varepsilon(\mathbf{x}, \mathbf{y}) \rightharpoonup p_t^{(0)}(\mathbf{x})$ | $\nabla \cdot \left( \bar{\kappa} \mathbf{K} \nabla p_t^{(0)} \right) = \frac{\bar{R}_{\mathbf{n}} \vert \Gamma_Y\vert }{\vert Y_t\vert } (p_t^{(0)} - p_c^{(0)})$ |
| | $\mathbf{u}_t^\varepsilon(\mathbf{x}, \mathbf{y}) \rightharpoonup \mathbf{u}_t^{(0)}(\mathbf{x}, \mathbf{y})$ | $\langle \mathbf{u}_t^{(0)} \rangle_{Y_t} = -\varepsilon \bar{\kappa} \mathbf{K} \nabla p_t^{(0)}$ |
| | $p_c^\varepsilon(\mathbf{x}, \mathbf{y}) \rightharpoonup p_c^{(0)}(\mathbf{x})$ | $\nabla \cdot \left( \mathbf{E} \nabla p_c^{(0)} \right) = 0$ |
| | $\mathbf{u}_c^\varepsilon(\mathbf{x}, \mathbf{y}) \rightharpoonup \mathbf{u}_c^{(0)}(\mathbf{x}, \mathbf{y})$ | $\langle \mathbf{u}_c^{(0)} \rangle_{Y_c} = -\mathbf{E} \nabla p_c^{(0)}$ |

For the sake of simplicity, from now on we denote by (*p*_*t*_, *p*_*c*_) the leading orders of the interstitial and capillary pressures 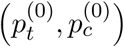.

### 3.1 Tensors properties

In order to ensure the well-posedness of the models that we have derived, the permeability tensors **K** and **E** need to be positive definite. This section is devoted to the analysis of the tensor properties with respect to the periodic cell *Y*.

#### Lemma 3.1. *The tensor* ***K*** *is symmetric and positive definite*.

*Proof*. Thanks to the Lax-Milgram theorem, problem (7) has a unique solution in *H*^1^(*Y*_*t*_)*/* ℝ. The variational formulation associated to (7) reads

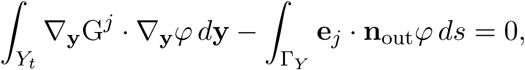

for any periodic *φ* ∈ *H*^1^(*Y*_*t*_) such that 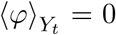. Considering *φ* = G^*i*^ on *Y*_*t*_, the following equations hold thanks to the divergence theorem

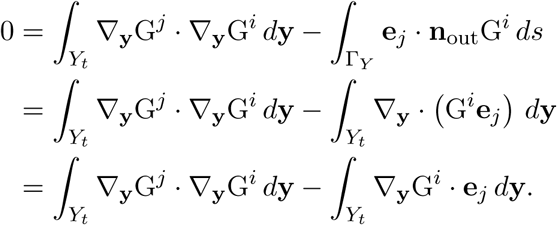

Therefore, the tensor **K** can be rewritten as

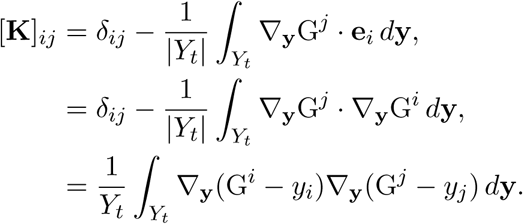

It follows that the tensor **K** is symmetric. To prove that the tensor is positive definite, we consider any ***λ*** ∈ ℝ^*N*^ and define

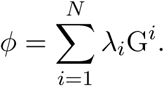

The function *φ* is periodic and belongs to the space *H*^1^(*Y*_*t*_). We prove that **K** is semi-positive definite:

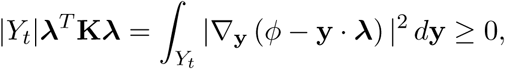

that is true for any ∇_**y**_ (*φ* − **y** · ***λ***). The equality holds if and only if

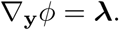

However, under the assumption of periodicity in a connected domain, ∇_**y**_*ϕ* = ***λ*** if and only if ∇_**y**_*ϕ* = ***λ*** = **0**. Therefore, **K** is positive definite.

#### Remark 3.2.

*The interstitial domain Y*_*t*_ *has to be connected to guarantee the positive definiteness of the tensor* ***K*** *(otherwise, it is semi-positive definite)*.

#### Lemma 3.3. *If the capillary domain Y*_*c*_ *is connected, then the tensor* ***E*** *is symmetric and positive definite*.

*Proof*. We proceed analogously as [40]. Thanks to the Lax-Milgram lemma, there exist a unique weak solution to problem (8), which variational formulation reads as

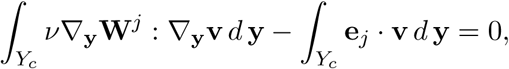

for any periodic **v** ∈ *H*^1^(*Y*_*c*_) such that ∇_**y**_ · **v** = 0 and **v** · **n** = 0 on Γ_*Y*_. Taking **v** = **W**^*i*^ the following identity holds:

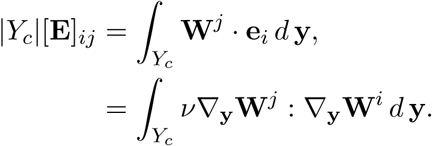

Therefore the tensor is symmetric. To prove that it is positive definite, we take any ***λ*** ∈ ℝ^*N*^ and define

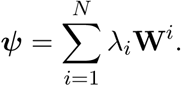

We first prove that ***λ***^*T*^ **E*λ*** is non-negative. Indeed,

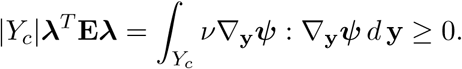

The equality holds if and only if ∇_**y**_***ψ*** = 0. Then, the following equation must be satisfied

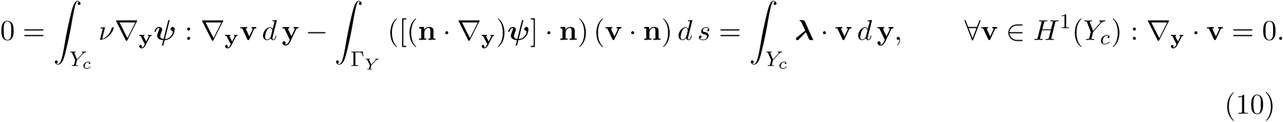

Since (10) holds for any **v** in the appropriate space defined above, it is valid also for **v** = ***λ***. Therefore, we conclude that (10) is true if and only if ***λ*** = **0** and state that **E** is positive definite.

#### Remark 3.4.

*When the domain Y*_*c*_ *is not connected, then the unique solution to problem* (8) *is* ***W***^*j*^ = ***0*** *and P*^*j*^ = *y*_*j*_. *In this case the tensor* ***E*** *is zero*.

#### Remark 3.5.

*The tensor* 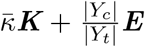 *is symmetric and positive definite since it is the sum of two symmetric and positive definite tensors*.

#### Remark 3.6.

*If one of the two domains* 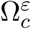 or 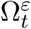 *is not connected, then p*_*c*_ = *p*_*t*_.

### 3.2 Limit problems

In this section, we show the link between the different models. We assume that both instertitium and capillary phases are connected so that the 3 models complemented with Dirichlet, Neumann or Robin conditions are well-posed.

### Equivalence of model 2 and 3

First consider model (2) and (3) with Dirichlet conditions. Assuming that both 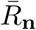 and *κ* are of order of magnitude *ε*, model (3) reads as follows:

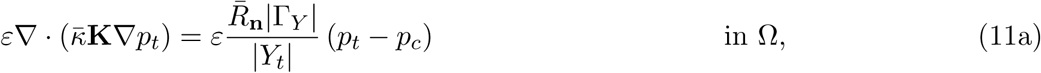

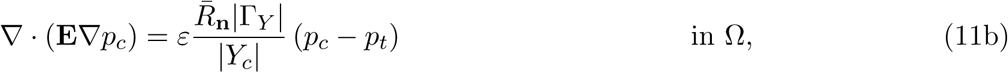

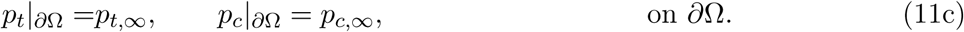

It is clear that (11b) is not a singluar perturbation of the operator ∇ · (**E**∇·) in the sense of Kato [42] and thus the solution to problem (11) tends to the solution to the following problem, which is nothing that model (3):

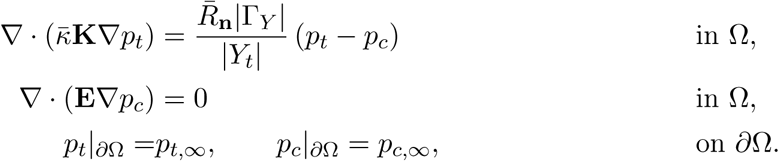

### Equivalence of model 1 and 2

Considering 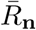 of the order of *ε*^−1^ and 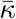 of the order of 1, model (2) of Table 1 reads then

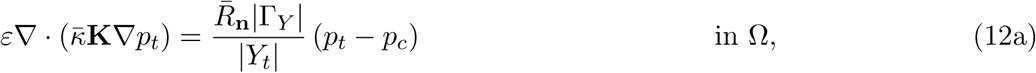

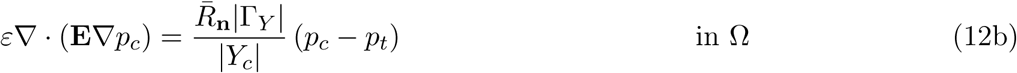

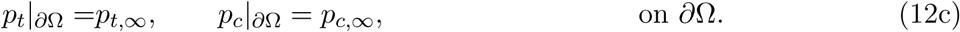

Here the asymptotic analysis is much trickier since both equations (12a)–(12b) are singular perturbation of the div-grad operator. In particular, a delicate asymptotic analysis makes appear a exponential decay of the *p*_*t*_ −*p*_*c*_ from the boundary, showing that out of the vicinity of the tumor boundary, both pressures are equal. The details of this results are given in [43], however we expose here the main arguments in the simple case where 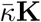 and **E** are colinear to the identity, that is for a *λ* ≠ 0:

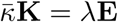

Then simple calculation shows that

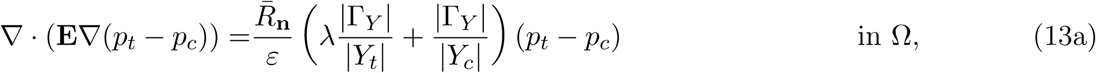

It is well-known, especially in conduction theory [44, 45] that problem (13) makes appear a so-called skin depth effect: the pressure difference *p*_*t*_ − *p*_*c*_ decays exponentially fast from the boundary. More precisely, denoting by *α* the factor given by

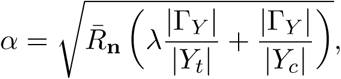

hence in the local coordinates near the boundary

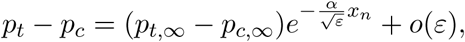

where *x*_*n*_ is the normal variable with respect to the tumor boundary.

Interestingly, we thus obtain that in this asymptotic regime, the solution to model (2) with Dirichlet boundary conditions can be approached by the solution to model (1) with the following appropriate boundary condition

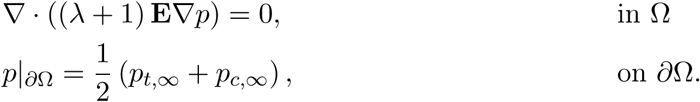

The rogorous proof is given in [43]. The result involves Riemannian geometry results which are far from the scope of this paper, however the general idea of the exponential decay of the pressure difference remains.

## 4 Numerical simulations

The Galerkin Finite Elements Method was used to discretize the equations in order to test the homogenized model in equation (2). 3D simulations were run in order to analyse the impact of the micro-scale geometry on the homogenized solutions and the influence of the vessel permeability *R*_**n**_ on the fluid transport. Regarding the homogenized models, the following strategy has been adopted:

- The periodic cell was considered as the unit cube (0, 1)^3^ in ℝ^3^. The domain was divided in two regions (*Y*_*t*_ and *Y*_*c*_) and the software Gmsh was used to perform the triangulation 𝒯_*h*_. Problem (8) was discretized with the Galerkin Finite Elements Method. Piecewise linear polynomials (𝕡_1_) were used for the variable P^*j*^. For the variable **W**^*j*^, we used piecewise linear polynomials with bubbles (𝕡_1*b*_ = {*v* ∈ *H*^1^(Ω) : ∀*K* ∈ 𝒯_*h*_ *v*|_*K*_ ∈ 𝕡_1_ ⊕ 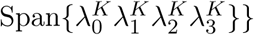, where 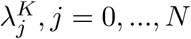, *j* = 0, …, *N* are the 4 barycentric coordinate functions of the element *K*). Problem (7) was solved on the domain *Y*_*t*_ using piecewise linear polynomials (𝕡_1_) for the variable G^*j*^.
- The tensors **K** and **E** were computed according to (9).
- The homogenized model was simulated on the normalized sphere of radius 0.5. Models in Table 1 were simulated using the Galerkin Finite Elements Method. Quadratic piecewise elements (𝕡_2_) were used for both *p*_*t*_ and *p*_*c*_.

### 4.1 Cell problems: tensor properties varying the microstructure

The tensors **K** and **E** defined in (9) have different properties according to the microstructure. To analyse them, we solved equations (7) and (8) in the unitary cell, i.e. the cube (0, 1)^3^ ⊂ ℝ^3^. Different geometric configurations for the domains *Y*_*t*_ and *Y*_*c*_ were tested (Figure 3).

**Figure 3.**
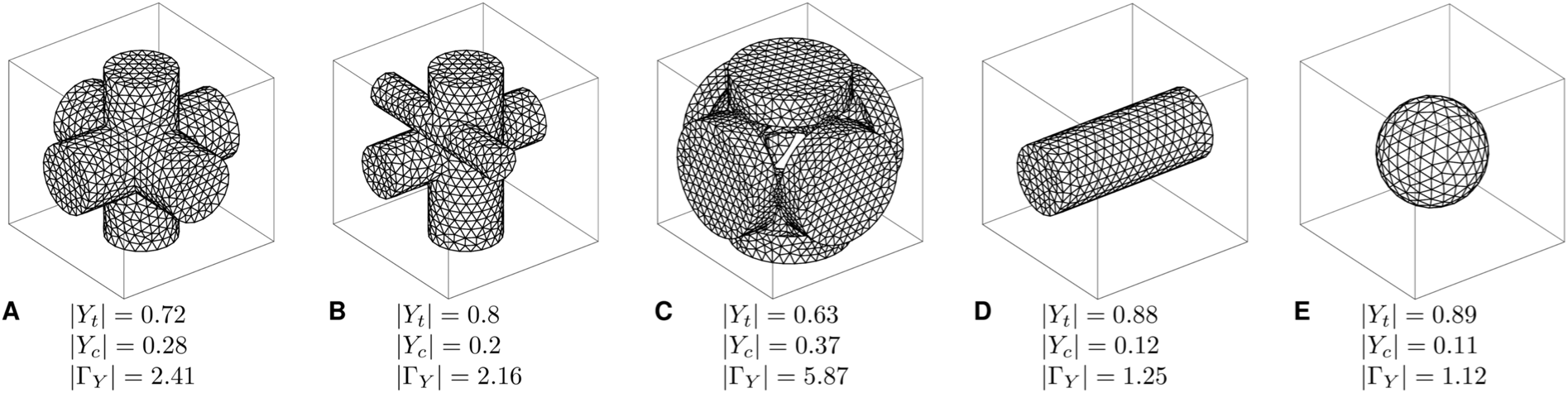
Different structures of the unit periodic cell with the respective volume and surface fractions. The mesh represents the capillary domain *Y*_*c*_, while the difference between the box and the mesh is the interstitial compartment *Y*_*t*_.

Table 2 provides the values of the elements in the two tensors **K** and **E**. These results confirm the analysis done in Section 3.1. Indeed, the tensors **K** and **E** are symmetric and positive definite when the two domains are connected (Fig. 3a, 3b and 3c). When the capillaries are not connected in all the directions (Fig. 3d, 3e), the tensor **E** is semi-positive definite as the solution to the cell problems (7) is trivial: **W**^*j*^ = 0 and P^*j*^ = **e**_*j*_, *j* = 1, 2, 3. Figure 3 provides the values of the interstitial and capillary volume fractions (|*Y*_*t*_| and |*Y*_*c*_|, respectively) and of the vascular surface Γ_*Y*_.

**Table 2.** Values of the tensors K and E for the different microstructures depicted in Fig. 3.

| | $\mathbf{K}_{11}$ | $\mathbf{K}_{12}$ | $\mathbf{K}_{13}$ | $\mathbf{K}_{21}$ | $\mathbf{K}_{22}$ | $\mathbf{K}_{23}$ | $\mathbf{K}_{31}$ | $\mathbf{K}_{32}$ | $\mathbf{K}_{33}$ |
| --- | --- | --- | --- | --- | --- | --- | --- | --- | --- |
| Fig 3a | 0.808 | 7.5e-5 | 7.89e-6 | 7.5e-5 | 0.808 | 5.49e-5 | 7.89e-6 | 5.49e-5 | 0.808 |
| Fig 3b | 0.877 | -1.76e-3 | 3.91e-3 | -1.76e-3 | 0.814 | 2.29e-3 | 3.91e-3 | 2.29e-3 | 0.933 |
| Fig 3c | 0.72 | -1.09e-4 | 1.03e-4 | -1.09e-4 | 0.72 | 3.98e-5 | 1.03e-4 | 3.98e-5 | 0.72 |
| Fig 3d | 1 | -3.19e-8 | -9.81e-8 | -3.19e-8 | 0.895 | 1.01e-4 | -9.81e-8 | 1.01e-4 | 0.895 |
| Fig 3e | 0.954 | -4.69e-5 | 5.2e-5 | -4.69e-5 | 0.954 | 8.14e-5 | 5.2e-5 | 8.14e-5 | 0.954 |
| | $\mathbf{E}_{11}$ | $\mathbf{E}_{12}$ | $\mathbf{E}_{13}$ | $\mathbf{E}_{21}$ | $\mathbf{E}_{22}$ | $\mathbf{E}_{23}$ | $\mathbf{E}_{31}$ | $\mathbf{E}_{32}$ | $\mathbf{E}_{33}$ |
| Fig 3a | 2.2e-3 | 8e-6 | -1.1e-6 | 8e-6 | 2.2e-3 | -1.3e-5 | -1.1e-6 | -1.3e-5 | 2.2e-3 |
| Fig 3b | 9.5e-4 | 8.1e-7 | 2.3e-5 | 8.1e-7 | 1.8e-4 | 7.7e-6 | 2.3e-5 | 7.7e-6 | 2.9e-3 |
| Fig 3c | 4.0e-4 | -8.7e-7 | -7.4e-7 | -8.7e-7 | 4.0e-4 | -2.7e-6 | -7.4e-7 | -2.7e-6 | 4.0e-4 |
| Fig 3d | 4.7e-3 | 0 | 0 | 0 | 0 | 0 | 0 | 0 | 0 |
| Fig 3e | 0 | 0 | 0 | 0 | 0 | 0 | 0 | 0 | 0 |

### 4.2 Macroscopic dynamic of fluid transport in tumours

We eventually considered realistic parameters to test model (2). The homogenized model was tested with a tumor considered as a sphere of normalized radius 0.5. Table 3 provides the values of the parameters of the model. Regarding the interstitial hydraulic conductivity *k*_*t*_, the vascular permeability *L*_*p*_ and the tumour characteristic length *L*, we considered values relative to different tissues, as summarized in Tables 4, 5 and 6, respectively. Simulations were run considering different microstructures, namely the ones shown in Figs 3a-c. Dirichlet boundary conditions were considered for the interstitial and capillary pressure, specifically *p*_*t*,∞_ = 0 and *p*_*c*,∞_ = 1 (normalized values).

**Table 3.** Fixed parameters used to simulate IFP and IFV.

| Parameter | Description | Value | Unit | Reference |
| --- | --- | --- | --- | --- |
| $\mu$ | blood viscosity | $4 \cdot 10^{-3}$ | $\text{kg} \cdot \text{m}^{-1} \cdot \text{s}^{-1}$ | [49] |
| $d$ | mean intercapillary distance | $50 \cdot 10^{-6}$ | m | [50] |
| $\alpha_{\text{BJ}}$ | BJS constant | 1 | - | - |
| $p_{t,\infty}$ | surrounding interstitial pressure | 0 | mmHg | - |
| $p_{c,\infty}$ | surrounding capillary pressure | [15,80] | mmHg | [7] |

**Table 4.** Values of the interstitial hydraulic conductivity *k*_*t*_ of different tissues.

| Tissue | $k_t$ [$\text{m}^3 \cdot \text{s} \cdot \text{kg}^{-1}$] | Reference |
| --- | --- | --- |
| Dog squamous cell tissue | $1.8 \cdot 10^{-12}$ | [51] |
| Mouse mammary carcinoma | $1.88 \cdot 10^{-13}$ | [52] |
| Hepatoma 5123 in vivo | $2.9 \cdot 10^{-15}$ | [53] |

**Table 5.** Values of the vessel permeability *L*_*p*_ of different tissues.

| Tissue | $L_p$ [$\text{m}^2 \cdot \text{s} \cdot \text{kg}^{-1}$] | Reference |
| --- | --- | --- |
| Mouse mammary carcinoma | $1.86 \cdot 10^{-10}$ | [52] |
| R3230 mammary adenocarcinoma | $4.5 \cdot 10^{-11}$ | [54] |
| Healty rat hindquarter tissue | $2.3 \cdot 10^{-12}$ | [55] |

**Table 6.** Characteristic length (diameter) of the tumour and corresponding tumour volume and value of *ε*.

| Characteristic length $L$ [mm] | Tumor volume [$\text{mm}^3$] | $\varepsilon = d/L$ |
| --- | --- | --- |
| 5 | 4.2 | 0.05 |
| 10 | 523.6 | 0.01 |
| 15 | 4200 | 0.005 |

#### Parameter influence

First, we looked at the behaviour of the solution varying the parameters *k*_*t*_, *L*_*p*_ and *L*. Examples of solutions as a function of the radius are shown in Fig. 4. In this case, we considered the microstructure of Fig. 3c. Results relative to the interstitial pressure and velocity were in agreement with the ones found in [2], where the authors considered the following model:

**Figure 4.**
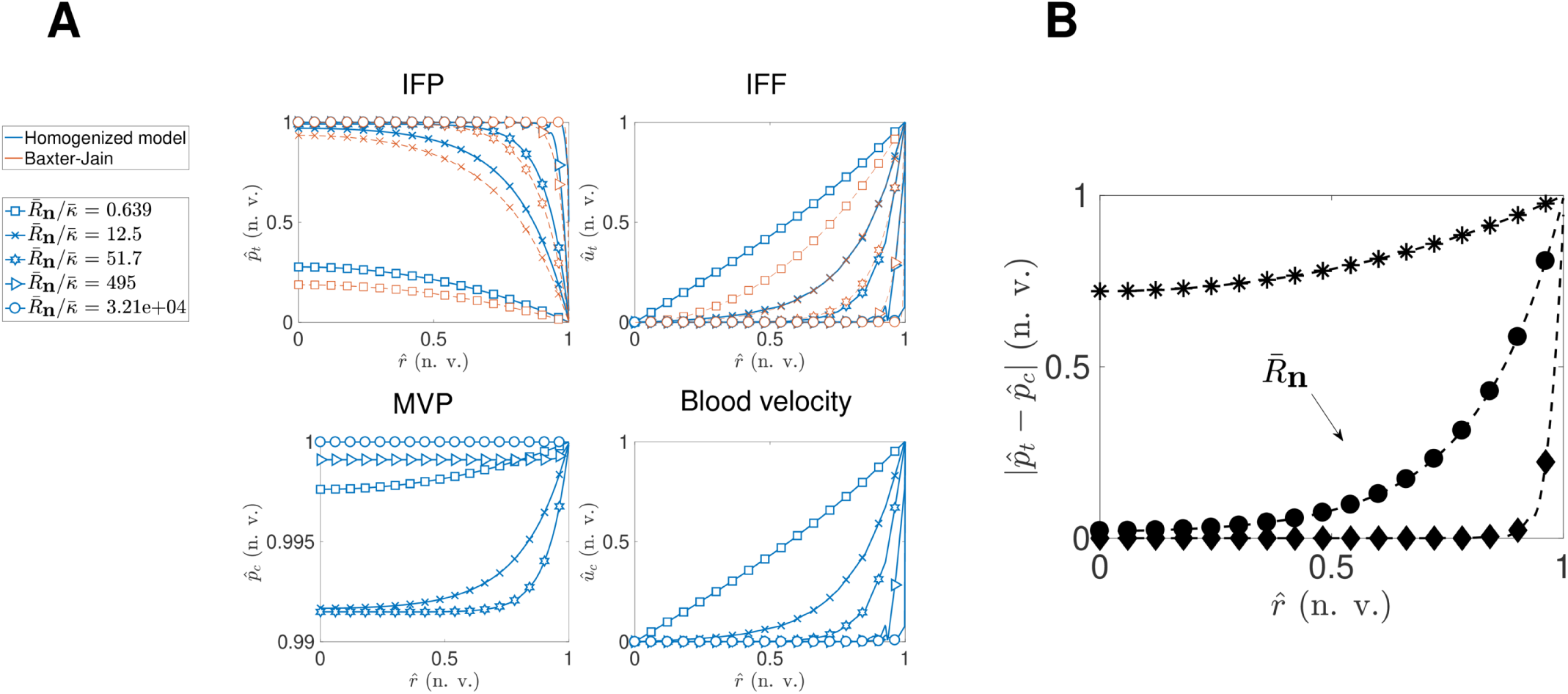
(A) Normalized values (n.v.) of interstitial fluid pressure and flow (IFP and IFF), of microvascular pressure (MVP) and of blood velocity as functions of the normalized radius 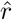 varying the parameter 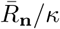. The microstructure considered in this case corresponds to Fig 3c. The blue lines are the simulations of the homogenized model (2) and the red lines are the results of Baxter and Jain model [2]. (B) Difference between 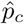 and 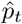 in normalized values (n.v.) as functions of the normalized radius 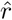 varying the parameter 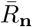 and with 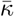 fixed.

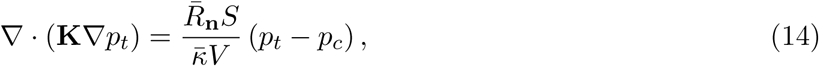

where the vascular pressure *p*_*c*_ is assumed to be constant and *S/V* is the vascular area per unit volume of the tumour. Therefore, we considered this value to be equal to |Γ_*Y*_|. The slight differences between the results obtained from the homogenized model and Baxter and Jain model (14) (Figure 4A) are due to the different rescaling of the equation, since we considered *S/V* to be the vascular area per unit volume of the interstitial compartment (|Γ_*Y*_ |*/*|*Y*_*t*_|).

The interstitial fluid pressure is large and almost constant in the centre of the tumour and has a sharp drop at the periphery for increasing values of 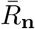 and decrasing values of 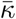. As a consequence, the interstitial fluid velocity is almost zero in the centre of the tumour (since the pressure gradient is close to zero) and large at the periphery. The microvessel fluid pressure is almost constant and close to the value at the boundary. For large values of the parameter 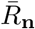, the capillary pressure decreases and gets closer to the interstitial fluid pressure. As a consequence, also the microvessel fluid velocity is close to zero in the centre of the tumour.

Eventually, we observed the skin depth effect of *p*_*c*_ − *p*_*t*_ when the permeability of the vessel walls increases (Fig 4B). Indeed, the pressure difference is almost zero at the centre of the tumour and increases exponentially in correspondence of the boundary.

#### Microstructure

We fixed the parameter values *k* _*t*_ = 1.8·1 0^−12^ m ^3^·s·k g^−1^, *L* _*p*_ = 1.86·1 0^−10^ m ^2^·s·k g^−1^ and *L* = 5 mm and looked at the behaviour of the solutions relative to the different microstructures. Fig. 5 shows the results relative to the unitary cells of Fig 3A-C. In all cases, the IFP shows a sharp drop at the periphery and it equates the capillary pressure in the centre of the tumour, while the capillary pressure is approximately constant in the whole tumour. The interstitial fluid velocity **u** _*t*_ is directed outward from the domain, while the blood velocity is directed inward. The two velocities are radially homogeneous in cases 5A and 5C, while they show asymmetries in case 5B due to the asymmetric microscopic structure of Fig 3B.

**Figure 5.**
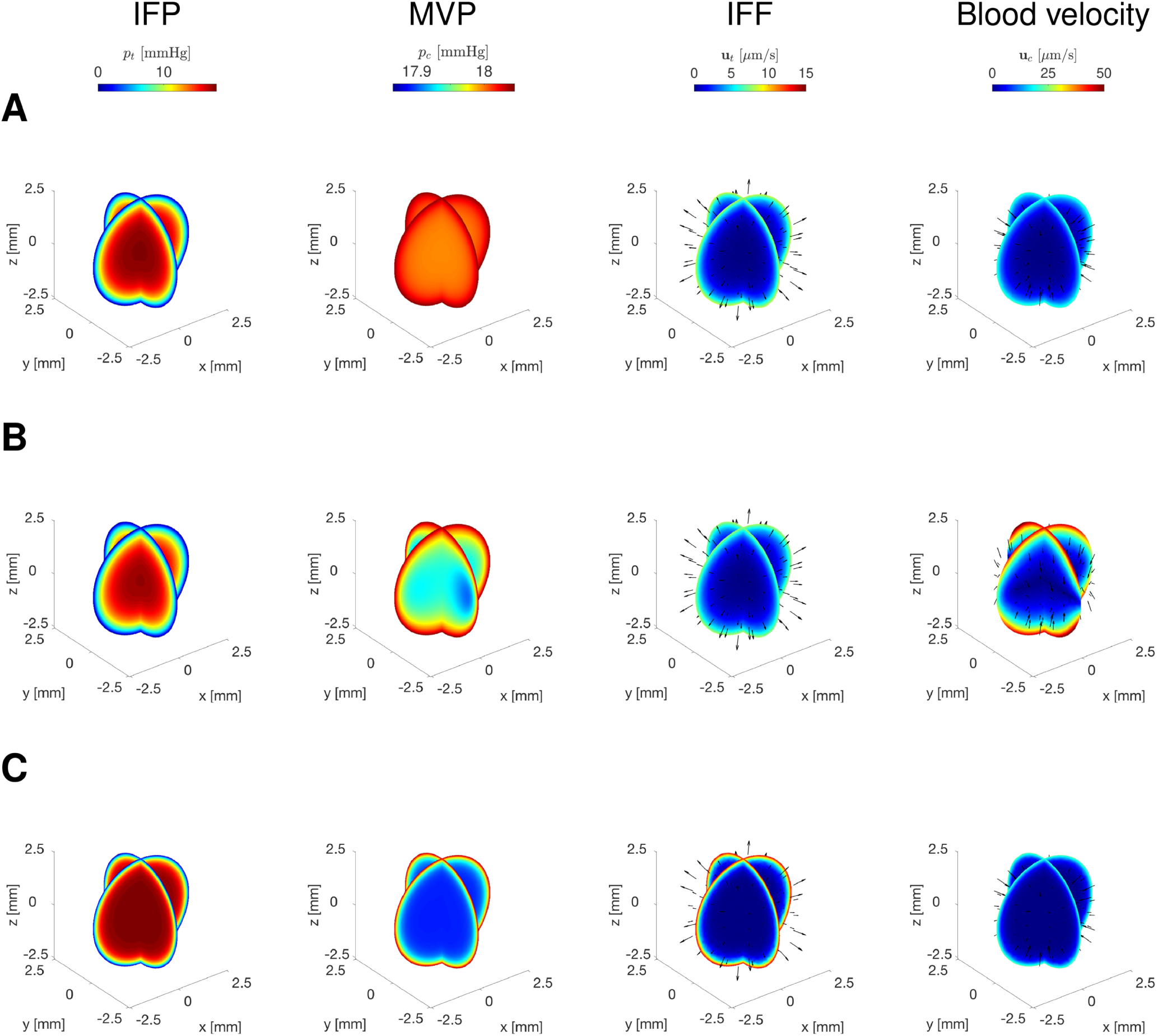
3D slices at the centre of the sphere with the interstitial pressure (first column), the capillary pressure (second comumn), interstitial velocity (third column) and capillary velocity (fourth column). Results were computed using the microstructure of Fig 3a (A), of Fig 3b (B) and of Fig 3c (C) and setting *k*_*t*_ = 1.8·10^−12^ m^3^·s·kg^−1^, *L*_*p*_ = 1.86·10^−10^ m^2^·s·kg^−1^ and *L* = 5 mm. IFP = interstitial fluid pressure, MVP = microvascular pressure, IFF = interstitial fluid flow.

We noticed that only when the capillary subdomain is smaller than the interstitial region, the blood velocity is larger than the interstitial fluid flow (data not shown). This is biologically relevant as the capillary volume fraction is usually within the range [16%, 50%] [46] and the average blood velocity is larger than the interstitial fluid velocity [47, 48].

#### Boundary conditions

Eventually, we tested model (2) with different boundary conditions. In particular, Neumann boundary conditions were considered for the capillary pressure, in order to ensure the continuity of the normal velocity in the vessels at the tumour periphery:

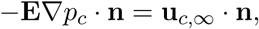

where **u**_*c*,∞_ is the blood velocity in the sourranding tissue. Dirichlet boundary conditions were imposed to the interstitial pressure. Well-posedness of model (2) is guaranteed with this set of boundary conditions 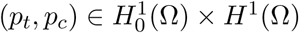.

We ran experiments with different boundary conditions for the capillary pressure *p*_*c*_ as summarized in Table 7. Homogeneous Dirichlet boundary conditions were considered for the interstitial fluid pressure *p*_*t*_. Figure S1 shows the results at the centre of the sphere as function of the normalized radius. The interstitial pressure increases at the centre of the tumour and equates the blood pressure in the three cases. When considering the case “Neumann 2”, the blood velocity is constantly high inside the domain and the capillary pressure profile is therefore due to the gradient along the *x*-axis.

**Table 7.** Different boundary conditions considered for the microvessel pressure *p*_*c*_.

| Experiment | Boundary condition (on $\partial\Omega$) | Parameter value (normalized) |
| --- | --- | --- |
| Dirichlet | $p_c = p_{c,\infty}$ | $p_{c,\infty} = 1$ |
| Neumann 1 | $-\mathbf{E}\nabla p_c \cdot \mathbf{n} = \mathbf{u}_{c,\infty} \cdot \mathbf{n}$ | $\mathbf{u}_{c,\infty} = -1 \cdot 10^{-3} \mathbf{n}$ |
| Neumann 2 | $-\mathbf{E}\nabla p_c \cdot \mathbf{n} = \mathbf{u}_{c,\infty} \cdot \mathbf{n}$ | $\mathbf{u}_{c,\infty} = [-1 \cdot 10^{-5}, 0, 0]^T$ |

## 5 Discussion

We have provided an analysis of the impact of microstructure properties of the tumour employing the homogenisation theory.

First, we have described a model at the microscopic scale that couples vascular, transvascular and interstitial fluids, adopting an asymptotic expansion technique. Then, we have derived three macro-scale models according to the vessel wall permeability and the interstitial hydraulic conductivity. After having analysed the well-posedness of the problems, we performed numerical simulations to assess some properties according to the microstructure.

Well-posedness is guaranteed when the two subdomains *Y*_*t*_ and *Y*_*c*_ are connected. When one region is not connected with respect to one axis, the fluid is not transported along this direction. For example, in Fig. 3e the capillary microstructure is a closed sphere, therefore there is no fluid transport in the blood vessels; in Fig. 3d, the vessel geometry is connected only along the *x*-axis that is the only direction for the capillary fluid flow. This represents a limit for the 2D simulations, as the subdomains *Y*_*t*_ and *Y*_*c*_ cannot be both connected. In this case, one among the interstitial or the vessel flow is always zero. However, tensors **K** and **E** can be determined by calibrating directly the homogenized models to medical imaging data.

Furthermore, we motivated the links between the various regimes and shown that model (2) covers a wide range of cases, confirming previous results [29]. In particular, we have shown that model (1) is equivalent to model (2) under certain conditions and that model (2) can be approximated to model (3) under certain assumptions on the parameters.

Eventually, we calibrated model (2) with parameters taken from the literature and analysed their influence on the solutions. We observed that different microstructures and different sets of boundary conditions strongly impact the macroscopic dynamics of the fluids. The geometric shape of the unitary cell influences the isotropy of the capillary fluid velocity, while the vascular volume fraction affects the blood velocity. Indeed, when the capillary volume fraction |*Y*_*c*_| is large, the blood velocity **u**_*c*_ is equal or lower than the interstitial fluid velocity **u**_*t*_. This might not be biologically relevant. On the other hand, when the capillary volume fraction is smaller the blood velocity is of higher magnitude and gets closer to the average values (around 1.62 mm · s^−1^ [56]). This confirms that the homogenized models are consistent with biological observations. Indeed, the vascular volume fraction lies within the values of 16% and 50%. [46, 57, 58]. Moreover, the average values of the pressures and of the velocities obtained from simulations with different sets of boundary conditions were compared against literature values. When Dirichlet-Dirichlet boundary conditions are considered, both the interstitial and the capillary pressures fit better the well-known profile of the IFP that is high at the centre of the tumour and shows a sharp drop at the periphery [2]. However, when Dirichlet-Neumann boundary conditions are considered for the interstitial and the capillary pressure, respectively, the blood velocity reaches average values closer to the literature ones. Possible improvements of our computations might be achieved by considering the correctors and by adding boundary layers, to take into account the Dirichlet boundary conditions that are imposed to the true solution 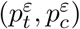 of the micro-scale model, but are not satisfied by the periodic solutions to the homogenized ones.

The current work focuses on the analysis of asymptotic models that describe fluid transport in tumour tissues. Fluid velocities are necessary to develop convection-diffusion models for the description of drug transport in tumour tissues. This motivated our choice of a steady-state model, as in reality, the time variation of the fluid transport is negligible with respect to the evolution of drug distribution inside the tumour. However, spatial tumour growth might be included in the model.

Further extensions might include a relaxation of the periodicity hypothesis, that might not be realistic in a biological context, as tumours are highly heterogeneous. Assuming a random distribution of the capillaries, it is possible to define properly the representative volume element to better predict the fluid flow in the tumour [59]. Moreover, rheological effects of blood should be included to model blood transport in capillaries [60].

Applications of the models include the incorporation of 3D imaging data. Images provide the microstructure of the vessel network, that is necesssary to compute the correctors.

## Supplementary Material

### S1: Derivation of the microscale model with asymptotic expansion

At the microscale, the domain Ω ∈ ℝ^*N*^ (with *N* = 2, 3) is the medium that consists of the interstitium Ω_*t*_, the vessel wall Ω_*m*_ and the capillary region Ω_*c*_. The interface between the capillary and the vessel wall and the one between the interstitium and the vessel wall are denoted respectively by Γ = ∂Ω_*c*_ ∩ ∂Ω_*m*_ and Γ_*δ*_ = ∂Ω_*t*_ ∩ ∂Ω_*m*_. Figure 1A shows the section of a capillary in the surrounding interstitium. In the three regions, the fluid flow is assumed to be incompressible. In the interstitium and in the capillaries, the equations explained in Section 2 hold.

Similarly to the interstitium, the capillary walls - of thickness *δ* - are considered as a porous medium with hydraulic conductivity *k*_*m*_, therefore the fluid flow velocity **u**_*m*_ and pressure *p*_*m*_ in the capillary walls are described by

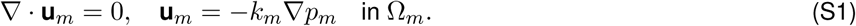

#### Interface conditions

At the two boundaries Γ and Γ_*δ*_, we have to consider interface conditions in order to couple the different equations. We make the following choices, similarly to [1]:

1. Continuity of the normal velocity on both Γ and Γ_*δ*_:

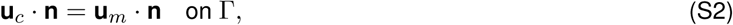

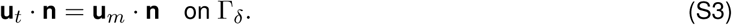 This condition guarantees the continuity of mass through the two interfaces and it is a natural choice since the fluid is assumed to be incompressible in the three regions.
2. Balance of the normal forces at the interfaces Γ, Γ_*δ*_:

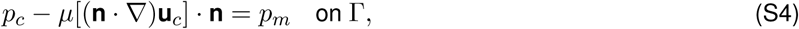

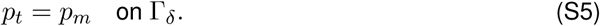 Condition (S4) is due to the fact that the blood force in Ω_*c*_ acting on Γ is equal to the normal component of the Cauchy stress vector [2], while the only force in Ω_*m*_ acting on the interface is the Darcy pressure *p*_*m*_. Analogously, equation (S5) is motivated by the fact that the only forces acting on the interface Γ_*δ*_ are the Darcy’s pressures *p*_*m*_ and *p*_*t*_ in the respective regions Ω_*m*_ and Ω_*t*_.
3. Beavers-Joseph-Saffmann condition on the tangential component of the capillary velocity at the boundary with a porous medium Γ:

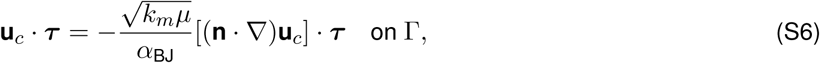

where *α*_BJ_ is a constant depending on the properties of the interface. This condition comes from the experimental evidence shown by Beavers and Joseph [3] who observed that the slip velocity along Γ was proportional to the shear stress along Γ. Equation of the form (S6) was derived by Saffmann using a statistical approach and the Brinkman approximation for non-homogeneous porous medium [4].

#### Non-dimensionalization

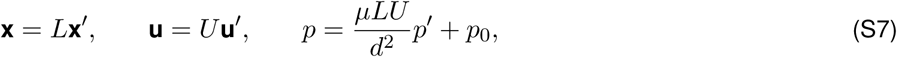

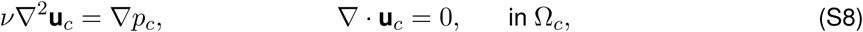

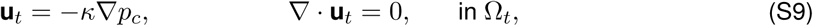

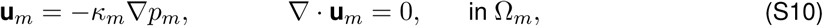

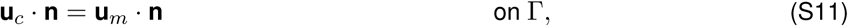

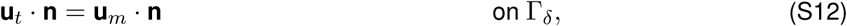

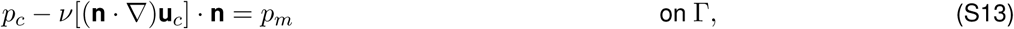

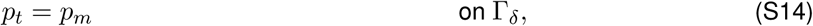

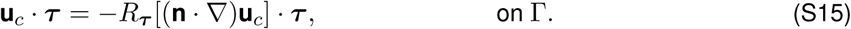

where

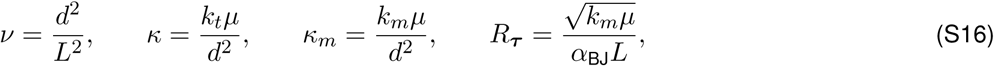

are dimensionless quantities.

#### Asymptotic expansion of the multi-scale model

We analyze the behaviour of the asymptotic system when the thickness of the capillary wall *δ* tends to 0, assuming that *κ*_*m*_ is proportional to *δ* with a proportionality coefficient *R*_**n**_ that will be defined later on:

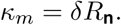

Let us denote by *η* the normal variable to the vessel membrane and by *θ* the tangential variable to the vessel wall. With these coordinates, the Laplacian is defined by

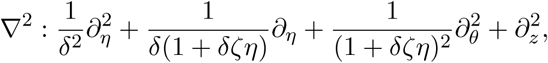

where *ζ* is the curvature of the section. Therefore, the fluid transport equations in the capillary wall and the interface conditions are given by

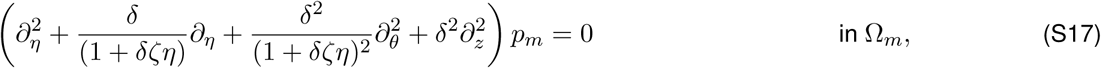

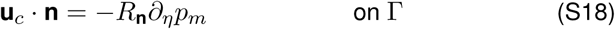

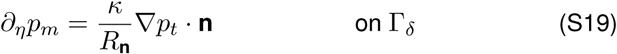

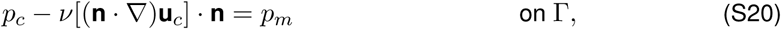

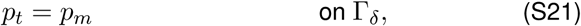

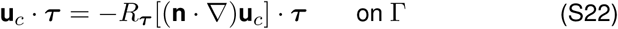

We perform an asymptotic expansion of the variables *p*_*m*_, *p*_*t*_, *p*_*c*_ and **u**_*c*_:

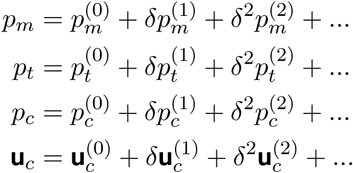

Equating coefficients of *δ*^0^ in (S17)-(S22), we obtain the following system of equations

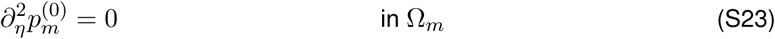

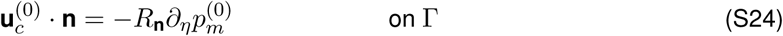

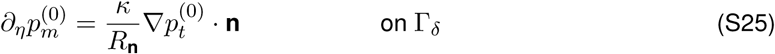

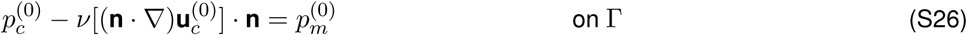

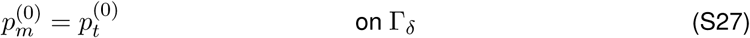

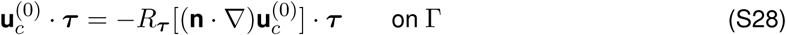

From (S23) and (S25) we infer

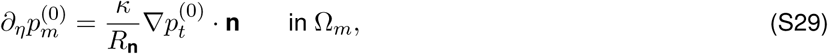

and then equations (S29) and (S26) leads to

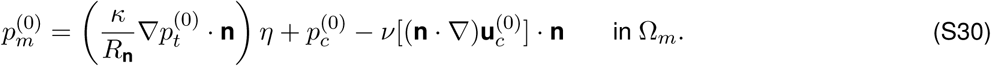

Therefore from (S27)

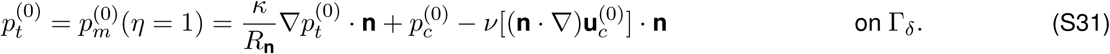

Let us define by Γ ∈ ℝ^*N*−1^ the boundary when *δ* goes to 0, i.e. when the two interfaces Γ_*δ*_ and Γ are superimposed. From equations (S31), (S24) and (S25) we derive formally the boundary conditions for small *δ*:

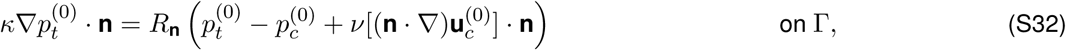

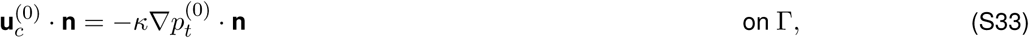

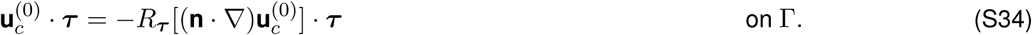

Conditions (S32)-(S33) can be rewritten as

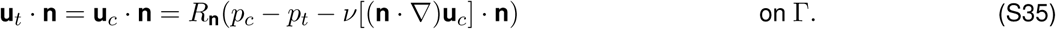

Equation (S35) is similar to Starling’s law, that is the most widely used equation in literature to model flux transport across the vessel wall [5, 6] and reads

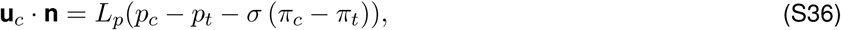

where *L*_*p*_ is the vascular permeability, *σ* is the osmotic reflection coefficient (*σ* ∈ (0, 1)) that expresses the glycocalyx filter function through the endothelial wall and (*π*_*c*_ − *π*_*t*_) is the oncotic pressure difference between the capillaries and the interstitium. However, the latter can be considered negligible compared to the interstitial fluid pressure difference in tumors [7, 8]. Moreover, the viscous term in equation (S35) is usually neglected but it is necessary to guarantee the well-posedness of the problem and does not change the physical meaning since it is based on the balance of the normal forces [9].

The relation between *R*_**n**_ and *L*_*p*_ is given thanks to the nondimensionalization of equation (S36):

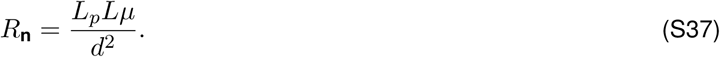

Therefore, our model is composed by equations (2), (2) in the respective regions of the domain Ω_*t*_ and Ω_*c*_ and by the interface conditions (S32)-(S34) on Γ.

### S2: Derivation of the macro-scale models

We derive the macroscopic models using formal two-scale homogenisation according to the magnitude of the permeability of the vessel wall and of the interstitial hydraulic conductivity, namely

- Model *γ*: = 0, *η* = 0;
- Model 2: *γ* = 1, *η* = 0;
- Model 3: *γ* = 2, *η* = 1.

The differential operators are then

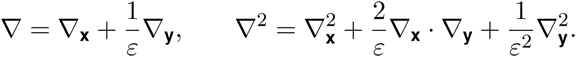

The fluid transport in the tumor tissue can be then written as follows:

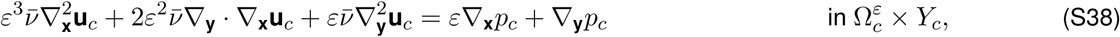

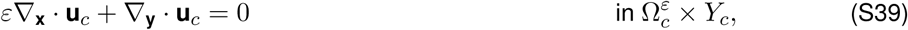

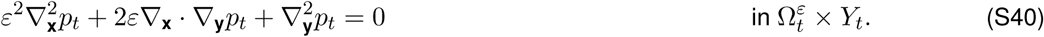

The interface conditions vary according to the value of*γ* and *η*:

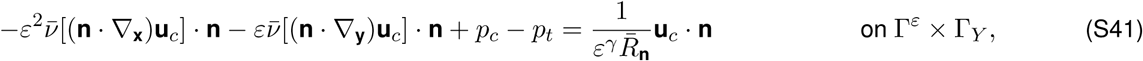

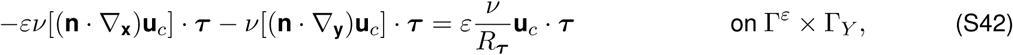

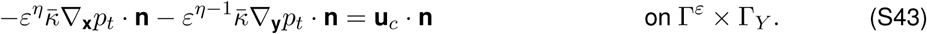

**Case 1: permeable vessel walls** *i*.*e. γ* = 0, *η* = 0.

When *γ* = 0, equation (S41) reads as

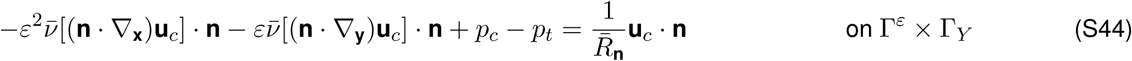

#### Identifying the terms of order *ε*^0^

In the interstitium, equating coefficients of *ε*^0^ in (S40) and (S43) gives

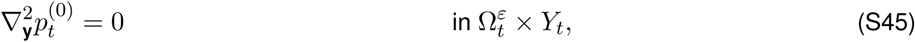

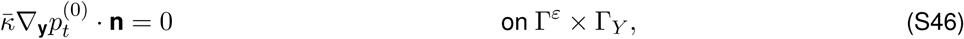

with 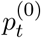 periodic in **y**. Therefore 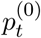 does not depend on the local scale, i.e. 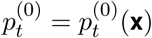.

Equating coeffiecients of *ε*^0^ in the capillaries gives:

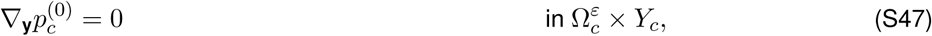

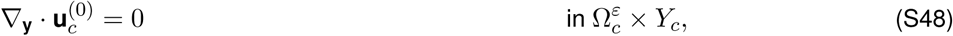

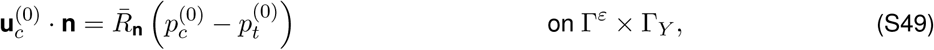

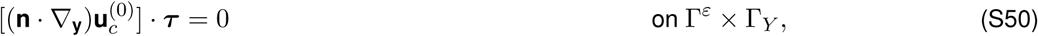

plus periodic boundary conditions on 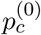 and 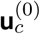 in **y**. Integrating equation (S48) we get the following condition on the pressure 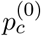:

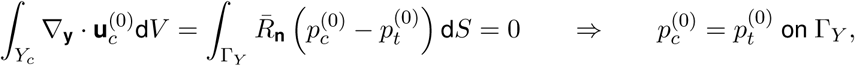

since 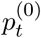 is a constant with respect to **y**. Therefore, 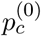 depends on the macroscale only and it is equal to 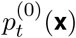 Moreover, condition (S49) becomes

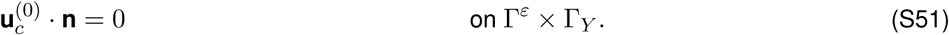

#### Identifying the terms of order *ε*^1^

Equating coefficients of *ε*^1^ in (S40) and (S43) yields

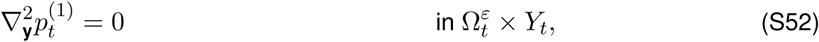

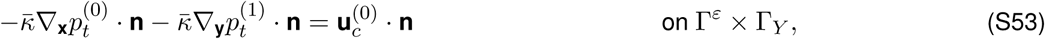

where 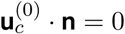. We exploit the linearity of system (S52)-(S53) and propose a solution of the form

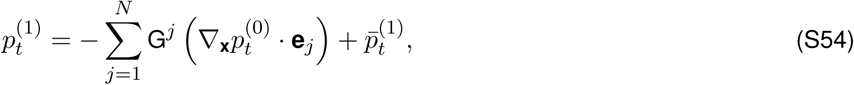

where G^*j*^ = G^*j*^ (**y**) satisfies the cell problem (7) for *j* = 1, …, *N*.

Equating coeffiecients of *ε*^1^ in the capillaries yields:

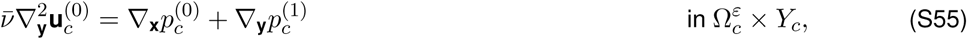

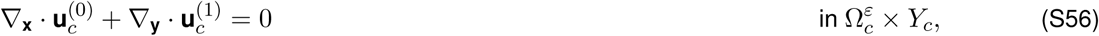

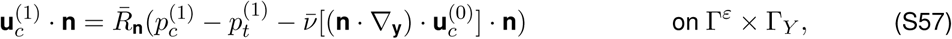

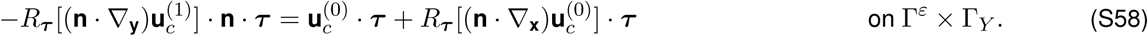

We exploit the linearity of the system composed by (S55)-(S48)-(S51)-(S50) and propose a solution of the form:

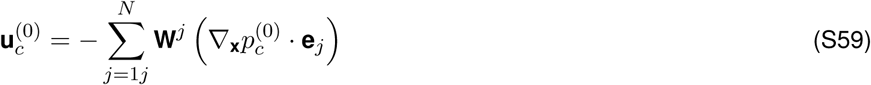

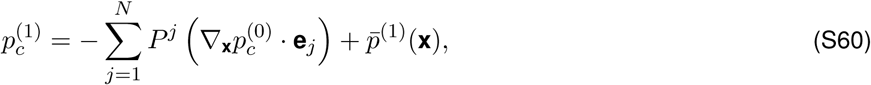

where (**W**^*j*^, *P*^*j*^) solve the cell problem (8) for *j* = 1, …*N*. Integrating (S59) over *Y*_*c*_, we find the leading order for the velocity 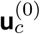:

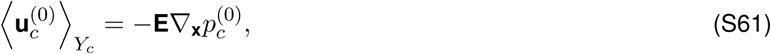

where **E** is defined in (9).

#### Identifying the terms of order *ε*^2^

Equating coefficients of *ε*^2^ in (S40) and (S43) gives:

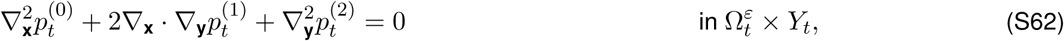

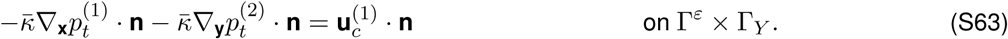

Integrating (S62) we obtain the equation for the leading order of the interstitial pressure:

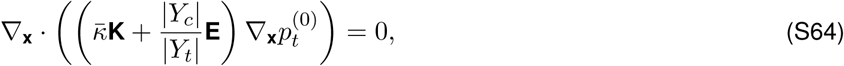

where **K** and **E** are defined in (9).

**Case 2: weakly permeable vessel walls** *i*.*e. γ* = 1, *η* = 0.

When *γ* = 1, equation (S41) becomes

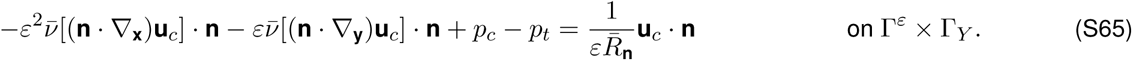

This case has been previously addressed [10, 9]. We write here the formal derivation of the macroscale model for the sake of completeness. Equations in the interstitium are the same as the ones in the previous case, while equations (S49) and (S57) take a different form in the capillaries. Nevertheless, the cell problems defined in (7) and (8) hold.

#### Identifying the terms of order *ε*^0^

As for the previous case, 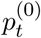 in the interstitium does not depend on the micro-scale, i.e. 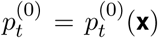. The interface conditions for the variables 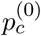 and 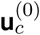 read as follows:

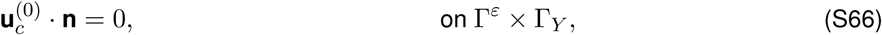

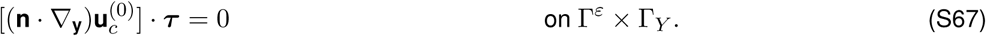

Equating coefficients of *ε*^0^ in the capillaries, equations (S47)-(S48)-(S66)-(S67) hold. Therefore, the leading order of the pressure in the capillaries depends only on the macro-scale 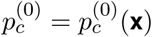.

#### Identifying the terms of order *ε*^1^

In the interstitium, the same results found in the previous case hold.

Equating coefficients of *ε*^1^ in the capillaries, yields

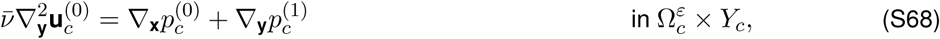

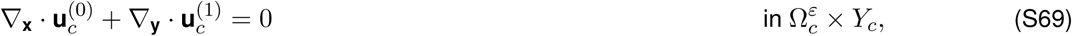

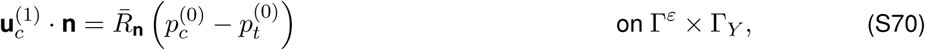

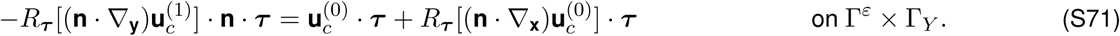

Exploiting the linearity of this system, we propose a solution of the same type of (S59) and (S60) where (**W**^*j*^, P^*j*^) solve the cell problem defined in (8).

#### Identifying the terms of order *ε*^2^

In the instersitial domain, equating coefficients of *ε*^2^ yields equations (S62) and (S63). We obtain the equations for the leading order by integrating (S59), (S70) in the capillaries, (S62) and (S63) in the interstitium:

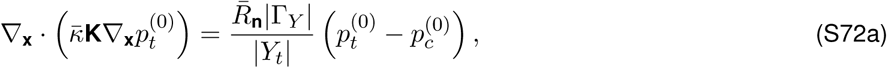

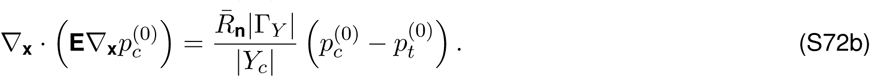

**Case 3: weakly permeable walls and weakly interstitial hydraulic connectivity** *i*.*e. γ* = 2, *η* = 1.

When *γ* = 2 and *η* = 1, equations (S41)-(S43) become

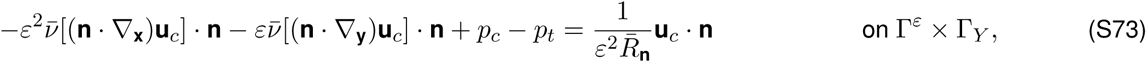

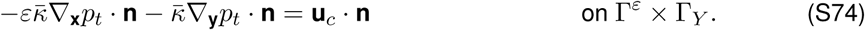

#### Identifying the terms of order *ε*^0^

Equating the coefficients of *ε*^(0)^, equation (S45) holds in the interstitium and (S47)-(S48) hold in the capillaries. The interface conditions for the variables 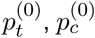 and 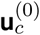 read as follows:

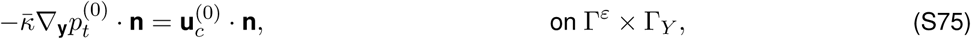

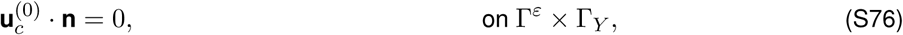

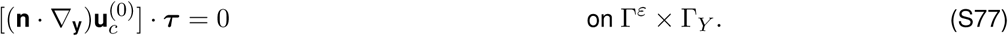

Since 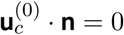 on 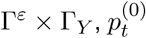 in the interstitium does not depend on the micro-scale, i.e. 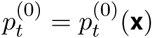.

Equating coefficients of *ε*^0^ in the capillaries, equations (S47)-(S48)-(S76)-(S77) hold. Therefore, the leading order of the pressure in the capillaries depends only on the macro-scale 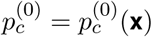).

#### Identifying the terms of order *ε*^1^

Equating coefficients of *ε*^1^, equation (S52) holds in the interstitium while equations (S55)-(S56) hold in the capillaries. The interface conditions for the variables 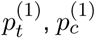 and 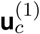 read as follows:

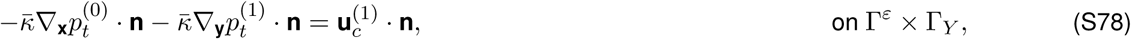

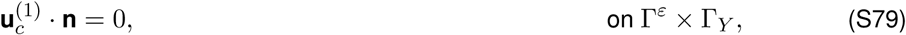

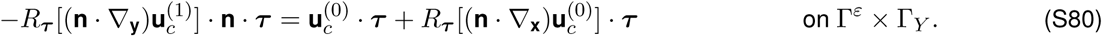

In the interstitium, since 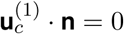 on Γ^*ε*^ *×* Γ_*Y*_, the same results for 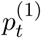 found in the previous cases hold.

In the capillaries, we exploit the linearity of the system (S55)-(S48)-(S76)-(S77) and propose a solution for 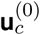 and 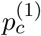 of the same type of (S59) and (S60) where (**W**^*j*^, P^*j*^) solve the cell problem defined in (8).

#### Identifying the terms of order *ε*^2^

Equating coefficients of *ε*^2^ in the interstitium, we obtain (S62) and the following interface condition:

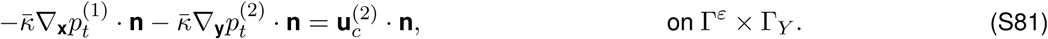

Equating coefficients of *ε*^2^ in (S42), the following interface condition holds:

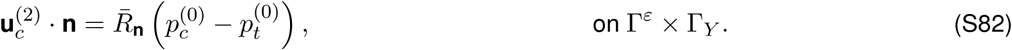

Integrating equations (S62) and (S81) we obtain the equation for the leading order of the interstitial pressure:

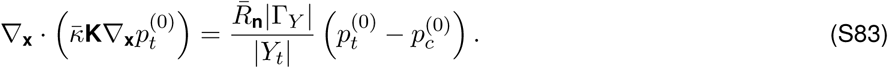

Integrating equation (S59) and (S79), we obtain the leading order for the pressure in the capillaries:

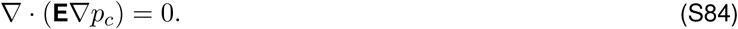

**Figure S1:**
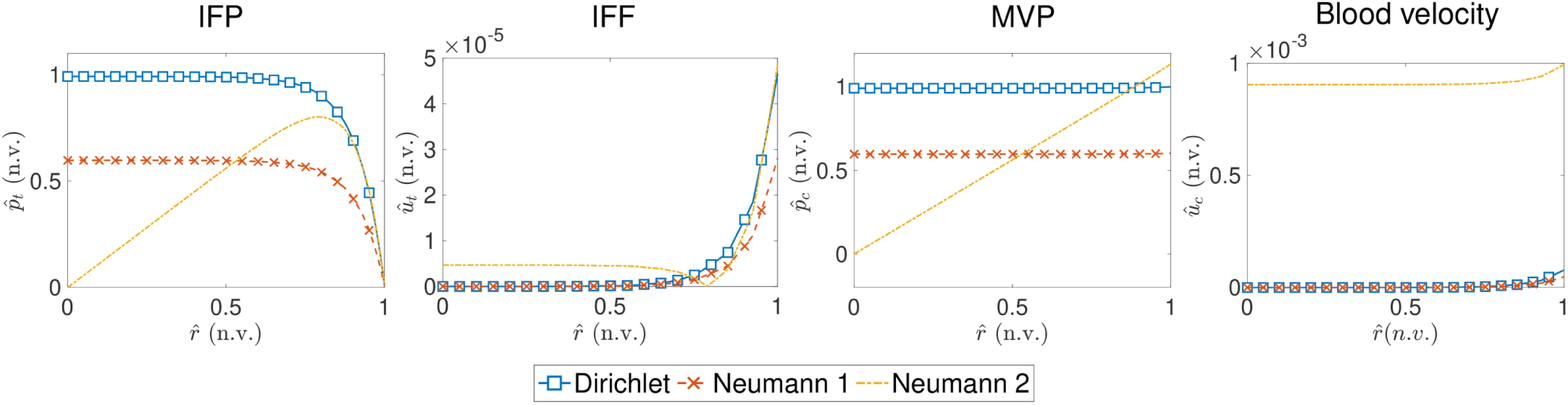
Normalized values (n.v.) of the interstitial fluid pressure (IFP) and flow (IFF), of the microvascular pressure (MVP) and of the blood velocity as functions of the normalized radius 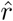 varying the boundary conditions. The microstructure considered in this case corresponds to Fig 3c.

## Notes

### Competing Interest Statement

The authors have declared no competing interest.

